# Structure and mutagenic analysis of the lipid II flippase MurJ from *Escherichia coli*

**DOI:** 10.1101/260596

**Authors:** Sanduo Zheng, Lok-To Sham, Frederick A. Rubino, Kelly Brock, William P. Robins, John J. Mekelanos, Debora S. Marks, Thomas G. Bernhardt, Andrew C. Kruse

**Affiliations:** Department of Biological Chemistry and Molecular Pharmacology, Harvard Medical School, Boston MA 02115, USA.; Department of Microbiology and Immunobiology, Harvard Medical School, Boston MA 02115, USA.; Department of Microbiology and Immunology, National University of Singapore, 117545, Singapore.; Department of Chemistry and Chemical Biology, Harvard University, Cambridge MA 02138.; Department of Systems Biology, Harvard Medical School, Boston MA 02115, USA.

## Abstract

The peptidoglycan cell wall provides an essential protective barrier in almost all bacteria, defining cellular morphology and conferring resistance to osmotic stress and other environmental hazards. The precursor to peptidoglycan, lipid II, is assembled on the inner leaflet of the plasma membrane. However, peptidoglycan polymerization occurs on the outer face of the plasma membrane, and lipid II must be flipped across the membrane by the MurJ protein prior to its use in peptidoglycan synthesis. Due to its central role in cell wall assembly, MurJ is of fundamental importance in microbial cell biology and is a prime target for novel antibiotic development. However, relatively little is known regarding the mechanisms of MurJ function, and structural data are only available for MurJ from the extremophile *Thermosipho africanus.* Here, we report the crystal structure of substrate-free MurJ from the Gram-negative model organism *Escherichia coli,* revealing an inward-open conformation. Taking advantage of the genetic tractability of *E. coli,* we performed high-throughput mutagenesis and next-generation sequencing to assess mutational tolerance at every amino acid in the protein, providing a detailed functional and structural map for the enzyme and identifying sites for inhibitor development. Finally, through the use of sequence co-evolution analysis we identify functionally important interactions in the outward-open state of the protein, supporting a rocker-switch model for lipid II transport.

## Introduction

The peptidoglycan (PG) cell wall is a defining feature of bacterial cell structure, and disrupting its synthesis is among the most effective strategies for treatment of bacterial infections. PG is synthesized from lipid II, which is first assembled on the inner leaflet of the plasma membrane and then flipped across the bilayer. The lipid II headgroup is then polymerized into PG by either class A penicillin binding proteins (PBPs) or SEDS proteins (1, 2). The size and hydrophilicity of the lipid II headgroup preclude spontaneous flipping across the membrane. MurJ is required for lipid II flippase activity *in vivo* (3). It is a member of the Multidrug/Oligosaccharidyl-lipid/Polysaccharide (MOP) family of transporters, which also includes proteins involved in similar biological processes such as the colanic acid flippase WzxC in bacteria and the proposed N-linked glycan flippase Rft1 found in all eukaryotes (4, 5).

Recently, the crystal structure of MurJ from the thermophilic bacterium *Thermosipho africanus* was reported (6). This high-resolution structure revealed an inward-open conformation not previously observed in any MOP transporter structure, and also confirmed that MurJ possesses two additional transmembrane helices at its carboxy terminus, which are not found in other MOP transporters. MurJ was crystallized in the presence of lipid II, and its structure showed a large central cavity containing electron density consistent with bound substrate. Although this crystal structure of *T. africanus* MurJ has been highly informative, this version of the protein is rather divergent from *Escherichia coli* MurJ (28% sequence identity) for which almost all functional studies have been performed to date. In order to better understand MurJ function, we crystallized *E. coli* MurJ by the lipidic cubic phase method, determined its structure to 3.5 Å resolution, and combined this structural analysis with high-throughput genetic and evolutionary covariation analysis.

## Methods

### Crystallography

A fusion protein was constructed, consisting of the crystallographic chaperone protein apocytochrome b562RIL (BRIL) fused to the amino-terminus of *E. coli* MurJ (residues 5 - 511), followed by a 3C protease site and a carboxy-terminal protein C epitope tag ("EDQVDPRLIDGK”). This was cloned into a pET28a expression vector using NcoI and NotI restriction enzymes (New England Biolabs), and the expression vector was transformed into *E. coli* BL21(DE3) strain. Liquid cultures were inoculated with transformed colonies and grown in LB medium at 37 °C with shaking. The cultures were shifted to 18 °C when the OD_600_ reached 0.8, and protein expression was induced at this point by addition of 0.5 mM isopropyl-β-D-thiogalactopyranoside (IPTG). After continued growth for 16 h, cells were harvested by centrifugation and resuspended in lysis buffer containing 25 mM HEPES pH 7.6 and 150 mM NaCl. After lysis by sonication, cell membranes were pelleted by ultracentrifugation at 35,000 rpm for 1 h. The pellets were homogenized using a glass dounce tissue grinder in a solubilization buffer containing 20 mM HEPES pH 7.5, 350 mM NaCl, 10% (v/v) glycerol, 2 mM CaCl_2_, 0.5% (w/v) lauryl maltose neopentyl glycol (LMNG; Anatrace), and 0.05% (w/v) cholesterol hemisuccinate (CHS; Steraloids). The sample was stirred for 1 h at 4 °C and clarified by centrifugation as before for 30 min. The supernatant filtered on a glass microfiber filter was loaded by gravity flow onto anti-protein C antibody affinity resin. The resin was washed extensively with buffer consisting of 20 mM HEPES, pH 7.6, 250 mM NaCl, 2 mM CaCl_2_, 0.01% LMNG and 0.001% CHS. The protein was eluted in the same buffer without CaCl_2_ supplemented with 5 mM EDTA and 0.1 mg/ml protein C peptide. The protein C tag was cleaved by 3C protease overnight. The protein was further purified by size exclusion chromatography on Superdex 200 increase in buffer 20 mM HEPES, pH 7.6, 250 mM NaCl, and 0.01% MNG and 0.001% CHS.

Prior to crystallization, the protein was concentrated to 45 mg/ml. Protein was reconstituted into lipidic cubic phase by the twin-syringe method (7), using a protein:lipid ratio of 1:1.5 (w:w). The host lipid was a 10:1 (w:w) mixture of monoolein (Hampton Research) with cholesterol (Sigma). Following reconstitution, protein samples were dispensed in 35 nl drops on glass sandwich plates using a Gryphon LCP robot (Art Robbins Instruments). Crystals were obtained with a precipitant solution of 100 mM MES pH 6.0, 100 mM potassium phosphate dibasic, 28% PEG 300, 1% 1,2,3-heptanetriol. Data collection was performed at Advanced Photon Source GM/CA beamline 23ID-B using a 10 μm beam diameter, 0.2 sec exposure, 5fold attenuation, and 0.2 degree oscillation angle. Data from 5 crystals were integrated, scaled, and merged in HKL2000 (8).

The crystals belonged to space group C222_1_ and contained one copy of BRIL-MurJ per asymmetric unit. Initial phasing was attempted by molecular replacement using *Phaser* (9), with recently published MurJ structure from *Thermosipho africanus* (PDB code 5T77) as a search model. This failed however, likely due to low sequence identity and large conformational differences between the template and model. Instead, we were able to solve the phase problem by *MR-Rosetta* in *Phenix* (10) with *T. africanus* MurJ as search model. This approach integrates molecular replacement, Rosetta modeling, autobuild and density modification and refinement (11). BRIL (PDB ID 4N6H) was subsequently placed using *Phaser.* The model was built in COOT and refined with *Phenix.* The final model incudes BRIL residues 3-107 followed by MurJ with residues 5-507. Ramachandran analysis showed that 95.05% of residues are in favorable regions and 4.95% are in allowed regions, with no outliers. Data collection and refinement statistics are summarized in **Supplemental Table 1**, and the structure is deposited in the Protein Data Bank under accession code 6CC4.

### Mutagenesis

Mut-seq was performed essentially as described before (1, 12). A plasmid library harboring mutagenized *murJ* (library size approximately 1 million) was transformed into *E. coli* strain CS7 (P_ara_::*murJ*) by electroporation and plated on LB with chloramphenicol and 0.2% (w/v) arabinose. Cells were scraped and the suspension was diluted to OD_600_ ≈1. After serial dilution to 10^−2^, 100 μl of the diluted suspension containing an estimated 10^6^ cells per mL was plated on LB chloramphenicol plates, supplemented with 100 μM IPTG, 0.2 (w/v) glucose, or 0.2% (w/v) arabinose. The plates were incubated overnight at 37 °C, and surviving colonies were harvested and diluted as described above. Plasmids were prepared from cell suspension before plating (input control), and after plating on LB with glucose, arabinose, or IPTG. Under this selection scheme, we do not detect any isolates without *murJ* insert on plates with glucose or IPTG as judged by colony PCR analysis of 16 isolates using GoTaq polymerase (Promega).

To prepare the sequencing libraries, purified plasmids from cell suspensions were digested with EcoRV and ClaI to release the cloned *murJ* inserts. The 2 kb fragments released were gel purified (Zymogen) and the amount of DNA recovered was quantitated using Qubit reactions (Invitrogen) according to the manufacturer’s protocol. After diluting the purified DNA samples to 2 ng/μl, 1 μl of the diluted DNA was mixed with 1.25 μl of TD buffer (Illumina 15027866) and 0.25 μl of TD enzyme (Illumina 15027865). The mixture was incubated for 10 min at 55 ^°^C in a thermocycler. Then, 11.2 μl of 2x KAPA master mix (KK2612 KAPA biosystems) and 4.4 μl of primers N705 and S503 (for the control input library), N706 and S503 (for the arabinose library), N701 and S504 (for the glucose library), and N702 and S504 (for the IPTG library). PCR was performed with 72 °C for 3 minutes, 98 °C for 5 minutes, 12 cycles of 98 °C for 10 seconds, 62 °C for 30 seconds, and 72 °C for 30 seconds, and a final extension step at 72 °C for 5 minutes. Quality of the PCR products was judged by running 3.75 μl of PCR product on a 1.5% agarose gel. The resulting products were purified and size selected by AMPure beads (Agencourt A63881). First, 18.75 μl of the PCR product was mixed with 15 μl of AMPure beads and incubated at room temperature for 5 minutes. The beads were captured by a magnetic stand, and the supernatant was discarded. The beads were then washed twice with 200 μl of 80% ethanol, air dried, and eluted by incubation with 40 μl of water for 5 minutes at room temperature. The eluted DNA solution was transferred to a new tube, and the size distribution of the libraries and the amount of DNA was measured by Tapestation (Agilent) and Qubit respectively. The prepared libraries were sequenced using Illumina Miseq v3 150 cycles kit (Illumina 15043893) exactly as described in the manufacturer’s protocol. In this case, 7 pM of the DNA was used and typically result in a clustering density around 900/mm^2^.

Data analysis was performed using CLC workbench (Qiagen). Failed reads were discarded and the sequences were trimmed with a cut-off quality value = 0.005 and a minimal length of 35 bp. The reads obtained were aligned using the global alignment setting to the reference sequence of pCS126 with the length and similar score set to 1. The unmapped reads from this alignment were collected, and further aligned to pCS126 with a similarity score set to 0.98, mismatch cost set to 2 and gap cost set to 3. Finally, the mutations from the reads were identified by the variant finder tool, using a setting of polyploidy = 1, minimal frequencies = 0.0001, and count = 2. The mutations found were then exported to Microsoft Excel for further analysis.

### Identifying evolutionary couplings in MurJ

The full-length sequence of MurJ (511 amino acids, MURJ_ECOLI) was used to generate multiple sequence alignments of different depths using 5 jackhmmer (13) iterations against the April 2017 Uniref100 database (14). To optimize both coverage of the query sequence and the number of sequences included in the alignment, all subsequent calculations were done on the alignment computed at bitscore 0.6 with 97.3% of the sequence including less than 30% gaps in each column and 15972 total sequences (4247 effective sequences after downweighting sequences with > 80% identity to the query). Evolutionary couplings (ECs) were then computed as described previously (15, 16), and a mixture model was applied to the set of all possible ECs to identify contacts that had a 99% probability of being in the tail of the score distribution (17). This threshold corresponds to 470 amino acid pairings, placing them in the 99.6^th^ percentile of the full space of potential couplings. These high-scoring ECs were then compared to the presented models, with a threshold of 5 Å for identifying true contacts.

## Results

Initially, we were able to obtain crystals for full length *E. coli* MurJ by the lipidic cupic phase method, but the crystals were small and diffraction was weak. We reasoned that the limited hydrophilic surface area of the protein for crystal packing may have prevented effective crystallization. To address this problem, we adopted a strategy from G protein-coupled receptor crystallography and fused the protein BRIL to the transporter amino terminus (18). The resulting protein crystallized readily, and crystals showed improved diffraction, enabling collection of a dataset to 3.5 Å resolution. The structure was subsequently solved using MR-Rosetta phasing with *T. africanus* MurJ as search the search model, and the structure was refined to R_work_/R_free_ of 0.28/0.30 with good geometry (**Supplemental Figure 1A, B** and **Supplemental Table 1**). As expected, the BRIL promotes tight crystal packing through interaction with itself from neighboring molecules along the **a** axis of unit cell (**Supplemental Figure 1C**). The residues 5-14 of TM1 which is supposed to be a straight helix is distorted due to its fusion to BRIL (**Figure 1**).

**Figure 1.**
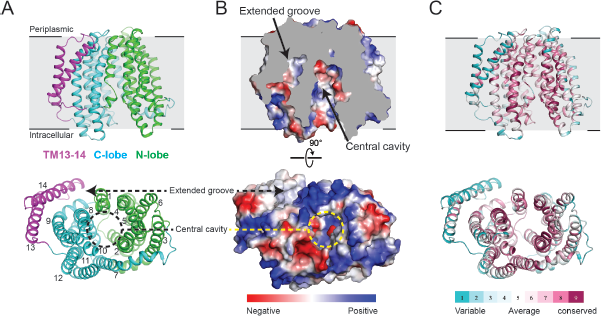
Overall structure of MurJ from *E. coli.* A, Ribbon representation of MurJ viewed from side and from the intracellular face, with N-lobe (TM1-6) and C-lobe (TM7-12) colored in green and cyan respectively, and TM13 and 14 in magenta. B, Electrostatic potential surface colored from red to blue for negatively to positively charged regions reveals a highly hydrophilic central cavity and hydrophobic extended groove formed by TM13 and TM14. The structure is shown in the same orientation as in panel A. C, The structure is colored by sequence conservation using ConSurf server with 483 MurJ sequences from both Gram-negative and positive bacteria, revealing a highly conserved central cavity in contrast to variable peripheral region.

The crystal structure of *E. coli* MurJ is similar overall to that of *T. africanus* MurJ, consisting of two homologous six-pass transmembrane bundles followed by a carboxy-terminal pair of helices located adjacent to the second bundle (**Figure 1**). The two six-helix bundles form two distinct lobes of the protein, which contact each other only on the periplasmic face of the protein, resulting in an overall inward open conformation similar to the structure of *T. africanus* MurJ. However, the cytosolic gap between the two lobes of the protein is roughly 4 Å smaller in the case of the *E. coli* protein (**Figure 2**). This may be a result of the fact that *T. africanus* MurJ was crystallized in the presence of substrate, while *E. coli* MurJ is substrate-free, allowing it to adopt a partially occluded conformation. Alternatively, it may reflect intrinsic differences between the MurJ proteins from the two species. A cross-section through the protein shows that the hydrophilic cavity extends almost entirely through the membrane, with a thin hydrophobic barrier where the two lobes of the protein interact on the extracellular face of the membrane (**Figure 2**). As in the case of *T. africanus* MurJ, the *E. coli* enzyme shows a large, deep groove formed by TMs 13 and 14, confirming that this feature is conserved across evolutionarily distant species. In addition, TMs 13 and 14 in *T. africanus* MurJ are shifted outward, perhaps as a result of substrate binding, creating a larger groove compared to *E. coli* (**Figure 2**). This is consistent with its proposed role as the interaction site for the lipid II isoprenoid tail, which must remain in the hydrophobic lipid bilayer throughout the flipping process.

**Figure 2.**
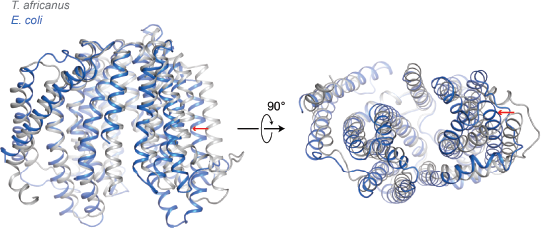
Structural overlay of MurJ from *T. africanus and E. coli.* The two structures were aligned at the C-lobe (left half of the protein as shown). *T. africanus* MurJ is colored in grey and *E. coli* in blue. *E. coli* MurJ reveals a relatively narrow central cavity and extended pocket compared to that in *T. africanus.* As a reference, the relative shift in orientation for TM3 is marked with a red arrow.

Many functionally important residues have been identified previously in *E. coli* MurJ, and our structure now provides a framework for their interpretation (19, 20). Of particular note is Asp39, which sits at the extracellular face of the protein and interacts with the positive dipole of TM8 and with Ser263 (**Figure 3**). This interaction bridges the two lobes of the protein, and may help to stabilize the inward open conformation. Asp39 is absolutely conserved among MurJ homologs in Gram-negative bacteria and is intolerant of substitution (19). Similarly, Ala29 sits at the interface between the two lobes of the protein, where it is almost entirely occluded. Mutation of this residue to Cys is known to be tolerated, but when labeled with a thiol-reactive agent the transporter exhibits complete loss of function (3). The steric enclosure of Ala29 would prevent labeling of this site in an inward-open state, but would be well accommodated in an outward-open conformation. Labeling of the A29C mutant would therefore trap the transporter in single conformation, accounting for the loss of lipid II flipping observed experimentally (3).

**Figure 3.**
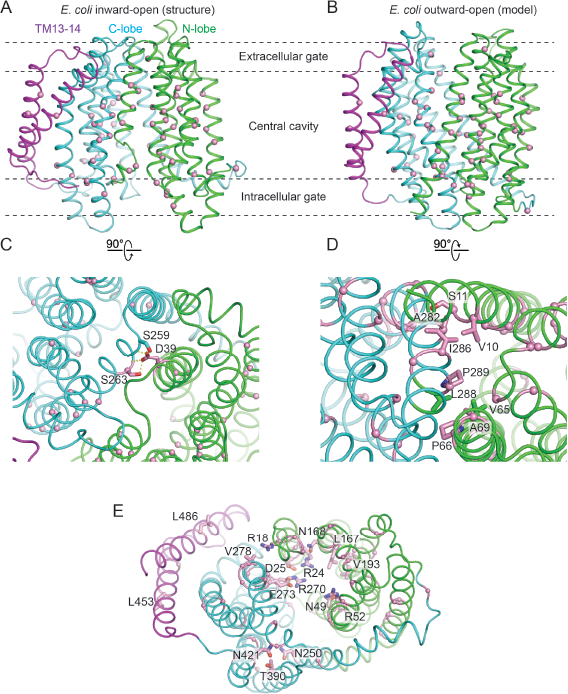
Mapping of Mut-Seq data on the MurJ structure. Residues at which mutations show more than 5-fold frequency decrease between cells grown in arabinose (MurJ_wt_ induction) and glucose (MurJ_wt_ depletion) are mapped in inward-open (A) and outward-open MurJ structure (B), shown as pink spheres in each case. C, Two essential residues shown in pink sticks mediate interactions between N-lobe and C-lobe in the extracellular side of inward-open MurJ. Dashed yellow line represents hydrogen bond. D, Several essential residues compose the intracellular gate of outward-open MurJ. A close-up view of hydrophobic interactions among these residues. E, Solvent-exposed essential residues located in the central cavity and extended groove (pink sticks).

To explore the functional consequences of MurJ mutagenesis in more detail, we took advantage of the genetic tractability and high transformability of *E. coli* to perform high throughput mutagenesis followed by next generation sequencing (“Mut-Seq”; (12)). This approach entails creating a large library of *murJ* mutants and then selecting for their ability to rescue loss of wild-type MurJ in cells. In brief, we constructed an *E. coli* strain in which wild-type *murJ* expression is driven by an arabinose promoter and a mutagenized plasmid-borne *murJ* by the *lac* promoter. In medium containing arabinose, wild-type MurJ is produced and the cells grow normally irrespective of whether or not the plasmid-borne *murJ* is functional. However, in glucose medium, wild-type *murJ* expression is repressed, and growth will rely on the expression of mutagenized *murJ* from the plasmid. Thus, only plasmids that encode functional *murJ* missense mutants will grow in this condition. By comparing the frequency of any given mutation between cell populations grown in the different media, a fitness measure of *murJ* mutations can be determined. Those changes that do not affect MurJ function are expected to be present at similar frequencies in the two populations, while those resulting in a loss of function are expected to be depleted from cells grown in glucose-containing medium.

Using this approach, we scored complementation of more than 1500 individual point mutants in *murJ,* including substitutions at all 521 codons in the sequence (**Supplemental Dataset**). Deleterious changes were identified throughout the protein coding sequence, including at known functional sites like Asp39 discussed above as well as many others. A total of 101 mutations resulted in at least a five-fold depletion in cells grown in glucose medium relative to cells grown in arabinose medium, suggesting a severely deleterious effect consistent with a complete loss of function. Indeed, 40 of these mutations result in the introduction of a premature stop codon. The other 61 mutations can be divided into four major groups: (i) Mutations in buried hydrophobic residues into charged or bulky residues, likely resulting in a loss of folding (**Figure 3A, B**); (ii) Mutations in residues composing the extracellular gate *(e.g.,* aforementioned D39, S263), located in the contact between two lobes in the extracellular side (**Figure 3C**); (iii) Mutations in solvent-exposed residues on the intracellular side, which likely comprise the intracellular gate and may stabilize the outward-open conformation. Mutations belonging to this fourth group are consistent with a MurJ homology model in the outward-open conformation, in which V65, P66 and A69 engage in extensive hydrophobic interactions with L288, P289 and S292 in the pseudosymmetry-related TM8 of the C lobe (**Figure 3D**). Mutations that insert large hydrophilic amino acids in these positions would disrupt the interaction between the two lobes, accounting for loss of function in these mutants. (iv) Mutations in solvent-exposed residues located in the central cavity and extended groove, which are likely involved directly in substrate binding or play a role in the regulation of transport (**Figure 3E**).

The function of MurJ in transport fundamentally depends on conformational change, but crystallography by nature presents only a static view of the protein. To gain insight into the possible existence of other conformational states, we turned to evolutionary coupling analysis. This technique can identify residues that have co-evolved with one another due to spatial proximity in the folded protein structure (16), and can reveal biologically important contacts irrespective of the structural conformation to which they belong. Using an alignment of 15972 sequences, we identified a total of 470 evolutionary couplings scoring in the 99^th^ percentile. The majority of co-evolving pairs correspond to residues that are located < 5 Å from one another in our structure (**Figure 4A, B**). However, a subset of strongly co-evolved residues are more distant from one another and do not make any direct contact. These co-varying pairs are located almost exclusively at the cytosolic face of the transporter, where they consist of pairs of residues on each of the two lobes of the protein (**Figure 4C, D**). To test whether the coevolution of these pairs can be accounted for by a conformational change to an outward-open state of MurJ, we constructed a outward-open homology model using the template of MATE from *Pyrococcus furiosus* (PDB code 3VVN) and TM13-14 from MurJ. When evolutionary couplings are superimposed on this model, cytosolic couplings are well accounted for while coevolving residues on the periplasmic face are now distant (**Figure 4B, D**). Taken together, these data show that both inward-open and outward-open states are under evolutionary selection pressure, providing strong support for a rocker-switch alternating access model of lipid transport.

**Figure 4.**
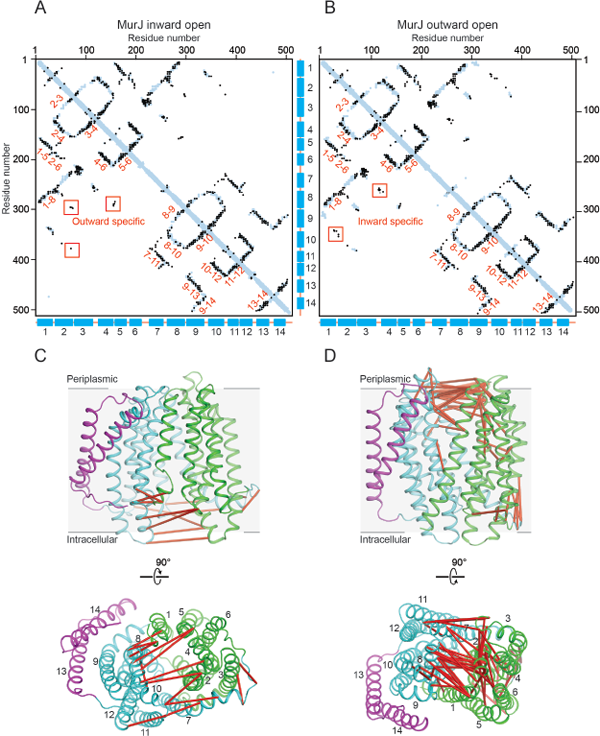
Evolutionary covariation analysis of MurJ. Residues pairs located less than 5 Å from one another in MurJ inward-open structure (A) and outward-open model (B) are shown as blue dots, while co-evolved pairs are shown as overlaid black dots. Strong covariations are observed between helices (labeled by helix number) which are adjacent to one another in the structure. A subset of strongly coevolved residue pairs are distant (>5 Å) in one conformation but close (<5 Å) in the other, indicating that both inward-open and outward-open states have been subjected to evolutionary selection. These violation pairs are mapped on the inward-open (C) and outward-open (D) model with red line connected.

## Discussion

Flipping of lipid II across the plasma membrane is an essential step in peptidoglycan biosynthesis, and represents a broadly conserved potential therapeutic target. Here, we have reported the crystal structure of *E. coli* MurJ, the most extensively studied of the broadly conserved MurJ transporters. While it is similar overall to a previously reported structure of *T. africanus* MurJ, the *E. coli* enzyme shows a distinct partially closed conformation, which may be representative of the apo-state of the protein. Moreover, the structure confirms that usual features seen in the *T. africanus* MurJ are conserved across species, attesting to their functional importance. Most notably, this includes a lateral gate between TM1 and TM8, and a long hydrophobic groove along TMs 13 and 14, which has been proposed as the binding site for the 55-carbon isoprenoid tail of lipid II. Consistent with a functionally important role for this region, high throughput mutagenesis experiments showed that introduction of premature stop codons between TM12 and TM13 is not tolerated (**Supplemental Dataset**). Similarly, several mutations in TM13 and TM14 (L453, L459 or L486) are depleted, further supporting a functionally important role for this region of the protein. In addition to TM13/14, high-throughput mutagenesis experiments highlighted other functionally critical regions of the protein including the inner (cytosolic) gate, the outer (periplasmic) gate, and the central cavity. Among poorly tolerated mutations, substitutions that altered the overall charge of the central cavity were particularly common, indicating the importance of electrostatic interactions for substrate recognition, transport, or both.

The identification of MurJ as the lipid II flippase is remarkably recent, and has been the subject of some controversy (21). In particular, other work had shown previously that the SEDS-family protein FtsW can flip lipid II *in vitro,* a result that is at odds with the identification of MurJ as the relevant flippase *in vivo* (22). However, *in vitro* flippase assays are notoriously difficult to conduct and interpret, in large part because the presence of improperly folded proteins can result in membrane defects that non-specifically scramble lipids (23). Recent data from a variety of studies has now solidified the identification of MurJ as the lipid II flippase, including mass spectrometry showing that lipid II binds strongly to MurJ but not the SEDS protein FtsW (24). Moreover, the SEDS proteins have now been unambiguously shown to catalyze peptidoglycan polymerization (1, 2), and it is difficult to imagine that the complex and biochemically disparate tasks of lipid II flipping and polymerization are mediated by a single enzyme. While SEDS proteins show sequence (and likely structural) similarity to glycosyltransferases, the structures of MurJ from *T. africanus* and *E. coli* show clear similarity to transporters. These proteins differ from other MOP-family transporters in exactly the ways expected for a lipid II flippase: they show a larger central cavity suitable to accommodate the bulky lipid II headgroup and they possess a lateral portal to allow the isoprenoid tail to remain in the lipid bilayer throughout flipping. In addition, we have shown that residues on opposite lobes of the cytosolic face of the protein show strong co-evolution, indicative of a transport cycle alternating between inward-open and outward-open states. Finally, the fact that homologous lipid flippases can substitute for MurJ in certain cases (25, 26)further supports its role as the lipid II flippase in cell wall assembly.

With the structure of *E. coli* MurJ, interpretation of the large body of mutagenesis and other functional data is now straightforward. Together with the previously reported structure of *T. africanus* MurJ, the structure and saturating mutagenesis of *E. coli* MurJ provides a framework for the design of future experiments to investigate unresolved aspects of MurJ function. These including identification of the energetic factor(s) driving lipid II flipping, and understanding how MurJ and other proteins in the peptidoglycan synthesis machinery work in concert to assemble the cell wall. In the long term, a detailed mechanistic understanding of cell wall assembly will not only be of value for understanding bacterial cell biology, but will also facilitate discovery of novel antibacterial agents.

## Author contributions

S.Z. performed MurJ purification, crystallization, and structure refinement with input from A.C.K. Mut-Seq experiments and data analysis were performed by L.-T.S. with supervision and input from W.P.R., J.J.M., and T.G.B. Evolutionary coupling analysis was performed by F.A.R. and K.B. with supervision from D.S.M. Overall project coordination was performed by A.C.K. with input from T.G.B. The manuscript was written by S.Z. and A.C.K. with input from all co-authors.

## Acknowledgments

This work was supported by CETR grant U19AI109764 (J.J.M., T.G.B., and A.C.K.), as well as R01GM106303 (D.S.M.). We thank Advanced Photon Source GM/CA beamline staff for excellent technical assistance with data collection. We thank all members of the Kruse and Bernhardt labs for helpful discussion and advice.

